# Think Like an Expert: Neural Alignment Predicts Understanding in Students Taking an Introduction to Computer Science Course

**DOI:** 10.1101/2020.05.05.079384

**Authors:** Meir Meshulam, Liat Hasenfratz, Hanna Hillman, Yun-Fei Liu, Mai Nguyen, Kenneth A. Norman, Uri Hasson

## Abstract

How do students understand and remember new information? Despite major advances in measuring human brain activity during and after educational experiences, it is unclear how learners internalize new content, especially in real-life and online settings. In this work, we introduce a neural measure for predicting and assessing learning outcomes. Our approach hinges on the idea that successful learning involves forming the “right” set of neural representations, which are captured in “canonical” activity patterns shared across individuals. Specifically, we hypothesized that understanding is mirrored in “neural alignment”: the degree to which an individual learner’s neural representations match those of experts, as well as those of other learners. We tested this hypothesis in a longitudinal functional MRI study that regularly scanned college students enrolled in an introduction to computer science course. We additionally scanned graduate student “experts” in computer science. We found that alignment among students successfully predicted overall performance in a final exam. Furthermore, within individual students, concepts that evoked better alignment with the experts and with their fellow students were better understood, revealing neural patterns associated with understanding specific concepts. These results provide support for a novel neural measure of concept understanding that can be used to assess and predict learning outcomes in real-life contexts.

## Introduction

Learning plays a central role in shaping our cognition. As we gain new knowledge, our thinking changes: as physicist Richard Feynman observed, “The world looks so different after learning science” (Feynman, 1969). Recently, multivariate “brain reading” analysis techniques have significantly advanced our understanding of how knowledge is represented in neuronal activity (Bauer and Just, 2019; Norman et al., 2006; O’Toole et al., 2007). These methods, together with representational similarity analysis (RSA), have made it possible to delineate the fine-grained structure of neural representations of learned knowledge, and to link neural patterns to specific knowledge across multiple domains (Bauer and Just, 2019; Haxby et al., 2014; Hsu et al., 2014; Mahon and Caramazza, 2011; Musz and Thompson-Schill, 2019; Parkinson et al., 2017). For the most part, this body of work has examined well-established concept representations rather than newly acquired concepts.

Recent imaging work has begun addressing this gap, examining the process of learning new concepts and extending a large body of work that has studied changes in neuronal circuits during and after learning (Karuza et al., 2014; McCandliss, 2010). Cetron et al. (2019) successfully used a multivariate neuroimaging approach to show that brain activity patterns recorded while students learned new categories in a Newtonian physics task can predict performance in a subsequent behavioral test. In an earlier study, Mason and Just (2015) reported a progression of activation throughout the cortex during learning, providing “snapshots” of the various cortical networks activated as participants progressed through explanations about different mechanical systems.

By design, these studies were conducted under carefully controlled experimental conditions. These required repeatedly exposing participants to a small set of static, discrete stimuli during a limited time, often within a single MRI scan. However, it is unclear whether results obtained under these conditions generalize to real-world settings. In a typical college course, students are required to communicate with instructors and peers; actively use a variety of static and dynamic learning resources inside and outside of class; assimilate multiple new concepts simultaneously; integrate course material over a prolonged period of several weeks; and often master new skills. Furthermore, selecting a course and taking it for credit is arguably different in terms of students’ motivation and interest than participating in a lab study. Therefore, a major goal of the current work was to examine learning in a real-life setting: a “flipped” introduction to computer science course in which students watched lecture videos outside of class.

A key part of learning is the communication of new information and its integration in students’ memory. Communication between individuals has been linked to neural “coupling”, such that (a) the brains of speakers and listeners show joint response patterns, and (b) more extensive speaker–listener neural coupling enables better communication (Hasson et al., 2012; Silbert et al., 2014; Stephens et al., 2010). Likewise, when people watch the same video, shared activity patterns emerge across the brain (Hasson et al., 2004). Recent imaging studies have shown that memories of this shared experience are encoded in a similar way across individuals, particularly in Default Mode Network (DMN) regions (Chen et al., 2017; Zadbood et al., 2017). Notably, specific concepts have also been shown to evoke similar neural activity patterns across individuals, suggesting a shared structure for neural representations (Bauer and Just, 2019; Mason and Just, 2016; Nguyen et al., 2019; Shinkareva et al., 2012; Yeshurun et al., 2017). This body of work suggests that shared neural responses reflect “thinking alike”. In the context of learning, the students’ goal could be viewed as laying the neural foundation that would allow them to think like experts.

Here, we sought to use shared neural activity patterns across learners and experts to quantify and predict understanding in a popular STEM course at Princeton University. We used class and expert patterns to model “canonical” representations and tested the hypothesis that alignment to these patterns reflected understanding. Our findings demonstrate that alignment during video lectures over the course of a semester successfully predicted final exam performance. We further found that, while verbally answering open exam questions, alignment between students and experts and alignment between students and classmates in medial cortical regions were both positively correlated with performance across questions, within individual students. Strikingly, a consistent set of relationships *between* topics emerged across students, correlating with performance within individual students and revealing how different concepts were integrated together. We thus show that approximating canonical neural representations supports understanding of STEM concepts learned in a college course setting. Our results provide support for a novel neural measure of concept understanding that is applicable in real life and online settings.

## Results

Did alignment to canonical neural representations emerge during learning, and did alignment reflect understanding of learned material? To address these questions, we examined neural activity patterns and learning outcomes in undergraduate students and in graduate “experts”. In collaboration with the Department of Computer Science at Princeton University, we recruited undergraduate students enrolled in COS 126: Computer Science - An Interdisciplinary Approach. The course introduces basic concepts in programming and computer science using a “flipped” classroom model, with lecture videos watched outside of class. Students underwent functional MRI (fMRI) scans five times during a 13-week semester while watching a subset of that week’s video lectures in the scanner. Subjects were asked not to view these lecture videos online before the scans. The subset of lectures shown in each scan was approximately 40 minutes long and comprised 3-5 segments (21 segments, 197 minutes in total). On the final week of the semester, students were shown - in the scanner - five 3-minute lecture recap videos with the highlights from previous weeks, followed by a final exam (Fig. 1a). To establish a baseline, the same exam was also given to students at the beginning of the semester, in written form. Graduate “experts” underwent the final scan only, watching the recap videos from all lectures and completing the final exam. The exam was self-paced, with exam questions (16 in total) spanning a variety of course topics from programming to theory. In the final exam, participants were asked to give verbal responses to visually presented questions (mean response length 31.9 seconds, s.d. 24.7). Questions were scored individually by course staff, providing a fine-grained measure of understanding.

**Figure 1.**
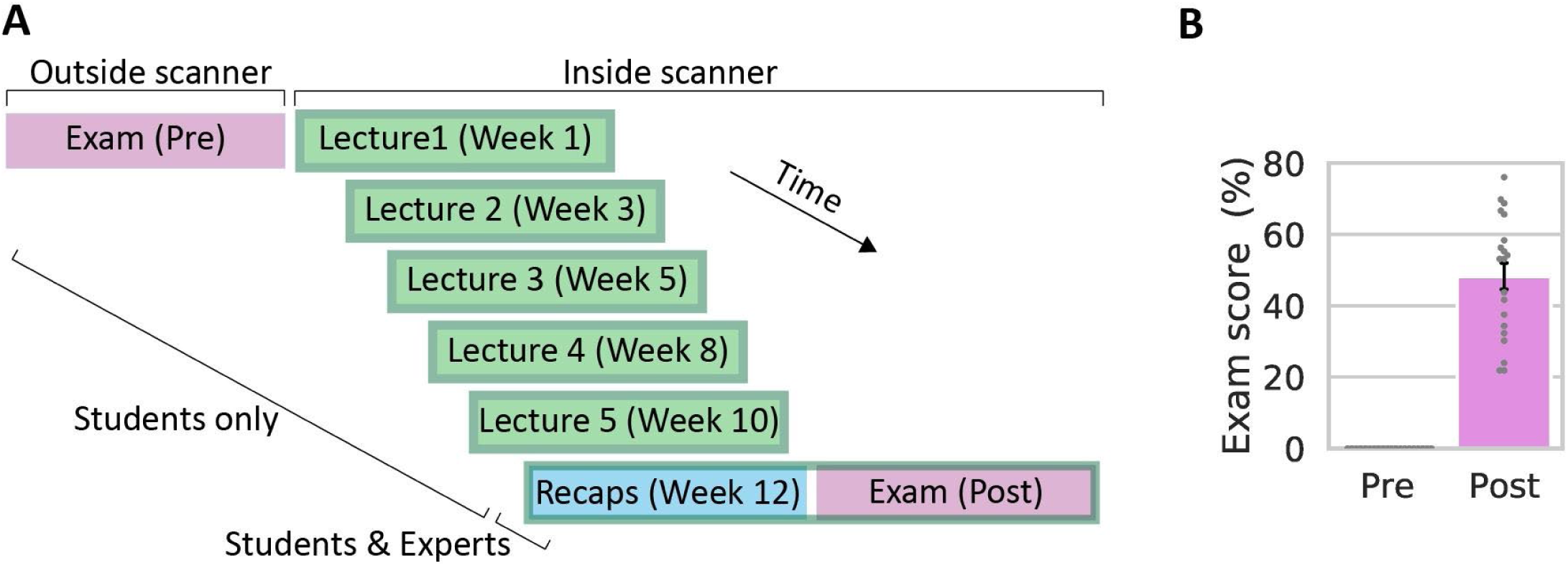
Study design and exam scores. **A**. Study design. Students enrolled in an introduction to computer science course underwent six fMRI scans throughout the course. During the first five scans, students were shown course lecture videos. On the final scan (bottom), students were shown lecture recaps and given a final exam. Experts underwent the final scan only. See table 1 for stimuli and task details. **B**. Exam scores. Pretest (left) was performed prior to scanning, posttest (right) was performed during scan 6. Individual students are shown in grey. Error bar, ±1 SEM.

All students received a score of zero on the baseline exam (Fig. 1b). This confirmed that students had no prior knowledge of course material. By the end of the course, all students demonstrated knowledge gains (two-sided t-test, t(19) = -12.6, p < 0.001), with substantial variance across students (range 22-76 out of 100, median 53, s.d. 17.1).

**Table 1.**
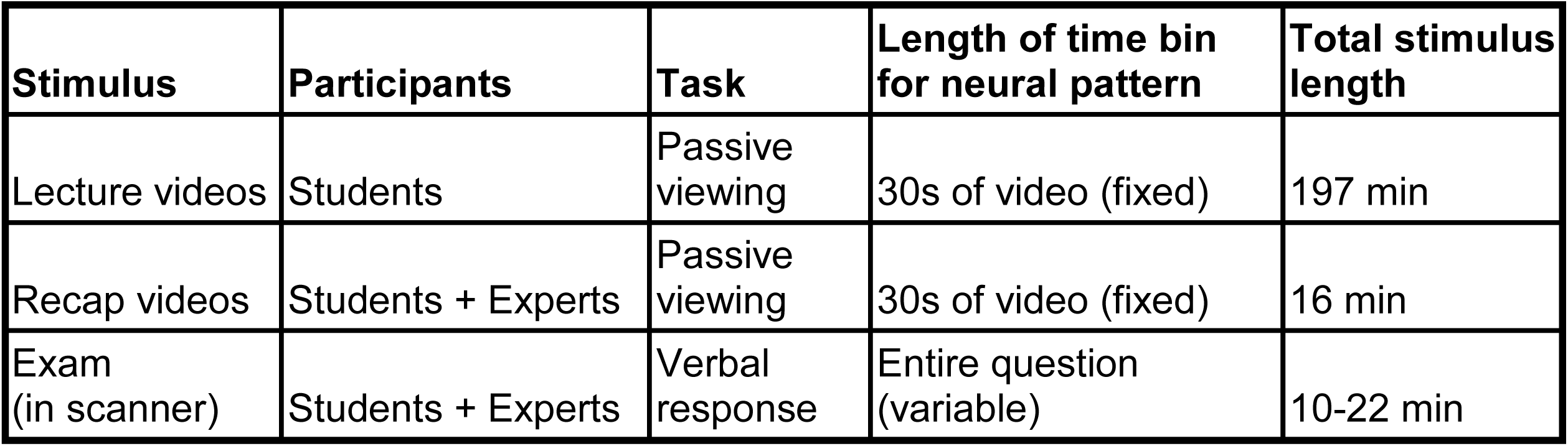
Stimuli and tasks.

### Prediction of learning outcomes from neural activity during lectures

Our first goal was to predict learning outcomes from brain activity during lecture videos. To this end, we calculated neural alignment-to-class across all lectures, comparing each student’s response patterns to the mean response patterns across all other students (Fig. 2A and Methods). Alignment values varied across the cortex, with the strongest values recorded in visual occipital regions, auditory and language regions, and parts of the default-mode and attention networks (Fig. 2B). The alignment map was in line with the body of literature showing that watching the same video elicits shared activity patterns across individuals (Hasson et al., 2004; Nastase et al., 2019). However, in the current work, alignment maps were not thresholded (i.e. statistical analysis for alignment effects was not performed), and all voxels were included in the subsequent searchlight analysis correlating alignment and exam scores. Correlation between alignment and exam scores was done using a between-participants design, first in eight anatomically-defined regions of interest (ROIs) in the DMN and hippocampus, and then across the entire cerebral cortex using a searchlight analysis. Our selection of ROIs was motivated by findings that activity in the DMN during memory encoding of new content (real-life stories or audio visual movies) predicted recall success for that material (Bird et al., 2015; Chen et al., 2017; Zadbood et al., 2017). The searchlight analysis enabled us to look for regions showing a correlation between alignment and learning outcomes in a data-driven manner. Throughout the manuscript, searchlight size was 5 × 5 × 5 voxels (15 × 15 × 15 mm cubes), and statistical significance evaluated using a one-sided permutation test (creating null distributions by shuffling labels 1000 times), controlling the false discovery rate (FDR) to correct for multiple comparisons at q = .05 (Benjamini and Hochberg, 1995; Kriegeskorte et al., 2008).

**Figure 2.**
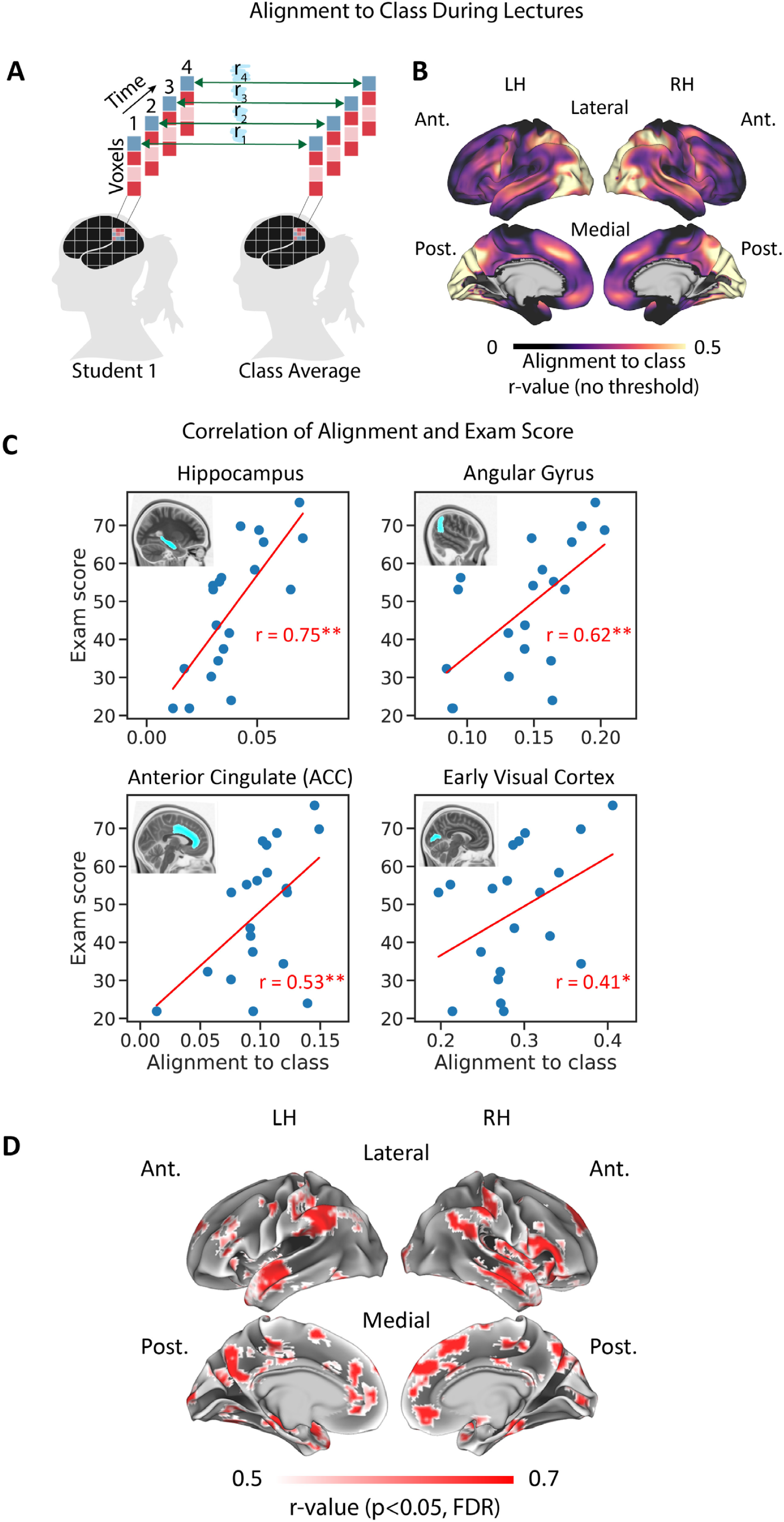
Alignment-to-class during lectures predicts final exam scores. **A.** Calculation of alignment-to-class during lecture videos. **B**. Alignment-to-class across the entire cerebral cortex. For demonstration purposes, this non-thresholded map shows mean alignment-to-class values across time and across students. **C**. Prediction of exam scores from neural alignment in example ROIs. Mean alignment-to-class across lectures (x-axis) is correlated exam scores (y-axis) using a between-participant design. Blue dots represent individual students. See table 2 for a summary of ROI results. **D**. Prediction of exam scores from neural alignment across the cortex. Searchlight analysis results shown. Voxels showing significant correlation are shown in color. LH, left hemisphere, RH, right hemisphere, Ant., anterior, Post., posterior.

Alignment-to-class in ROIs during lecture videos showed a significant positive correlation with final exam scores in the angular gyrus, precuneus, anterior cingulate cortex (ACC) (all overlap with the DMN), and the hippocampus, as well as early visual and auditory areas (Fig. 2C-D, Table 2). Across ROIs, the highest correlation values were observed in the hippocampus, allowing the most reliable prediction of learning outcomes. Our cortical searchlight analysis showed multiple brain regions where students’ alignment-to-class predicted their final exam scores (Fig. 2D). In line with the ROI analysis results, these regions included anterior and posterior medial areas as well as the bilateral angular gyrus, key nodes of the DMN. In addition, we observed significant correlations in temporal and insular cortex. A power analysis revealed that prediction improved as more data was aggregated across lectures (Fig. S1 and Supplementary Information).

**Table 2.**
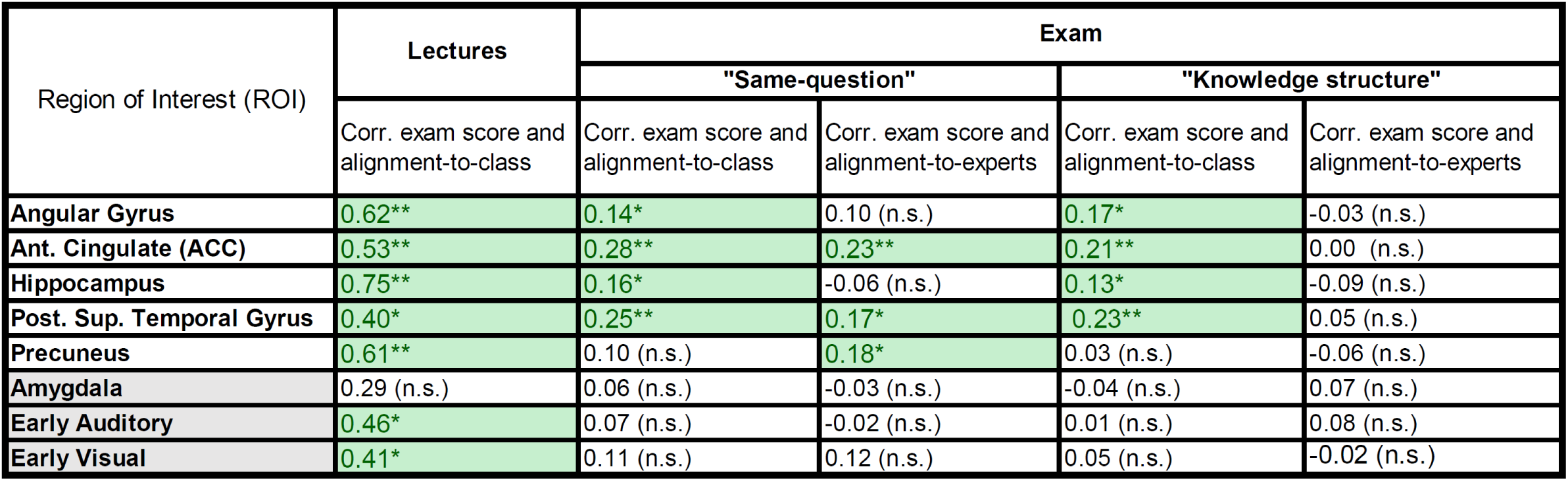
Prediction of exam scores from neural alignment in ROIs. Correlation between alignment measures and exam score during lectures and during the final exam. Results are shown in DMN ROIs as well as in control regions in sensory cortex (visual, intracalcarine cortex; auditory, Heschl’s gyrus) and subcortex (amygdala). Green, significant correlation (permutation test, p<0.05, FDR corrected across ROIs). n.s., not significant.

| Region of Interest (ROI) | Lectures | Exam |  |  |  |
| --- | --- | --- | --- | --- | --- |
|  |  | "Same-question" |  | "Knowledge structure" |  |
|  | Corr. exam score and alignment-to-class | Corr. exam score and alignment-to-class | Corr. exam score and alignment-to-experts | Corr. exam score and alignment-to-class | Corr. exam score and alignment-to-experts |
| Angular Gyrus | 0.62** | 0.14* | 0.10 (n.s.) | 0.17* | -0.03 (n.s.) |
| Ant. Cingulate (ACC) | 0.53** | 0.28** | 0.23** | 0.21** | 0.00 (n.s.) |
| Hippocampus | 0.75** | 0.16* | -0.06 (n.s.) | 0.13* | -0.09 (n.s.) |
| Post. Sup. Temporal Gyrus | 0.40* | 0.25** | 0.17* | 0.23** | 0.05 (n.s.) |
| Precuneus | 0.61** | 0.10 (n.s.) | 0.18* | 0.03 (n.s.) | -0.06 (n.s.) |
| Amygdala | 0.29 (n.s.) | 0.06 (n.s.) | -0.03 (n.s.) | -0.04 (n.s.) | 0.07 (n.s.) |
| Early Auditory | 0.46* | 0.07 (n.s.) | -0.02 (n.s.) | 0.01 (n.s.) | 0.08 (n.s.) |
| Early Visual | 0.41* | 0.11 (n.s.) | 0.12 (n.s.) | 0.05 (n.s.) | -0.02 (n.s.) |

### Neural alignment between students and experts

Neural alignment-to-class was strongly correlated with alignment-to-experts. Experts were scanned during recap videos (16 minutes in total) and while taking the final exam. We separately calculated alignment-to-class and alignment-to-experts for each student in each task and then correlated these measures using a between-participants design (see Methods). Figure 3A shows the results of this analysis in an example ROI in anterior cingulate cortex (ACC) during recaps, while Fig. 3B shows results in the same ROI during the exam. In both tasks, alignment-to-class and alignment-to-experts were positively correlated across all ROIs (Table 3). A searchlight analysis revealed that these effects extended to large parts of cortex, including the default-mode and attention networks. Cortical maps for recaps and the exam are shown in Fig. 3C and 3D respectively. These results indicate that the mean responses across all students converge to the average, or “canonical”, responses seen in experts during both recaps and the final exam. Furthermore, it indicates that the individual differences seen across subjects in their ability to be aligned to *class* are preserved when we look at their ability to converge to canonical *expert* responses. Next, we asked whether the ability of each student to converge to these canonical responses predicted learning outcomes during the final exam.

**Table 3.**
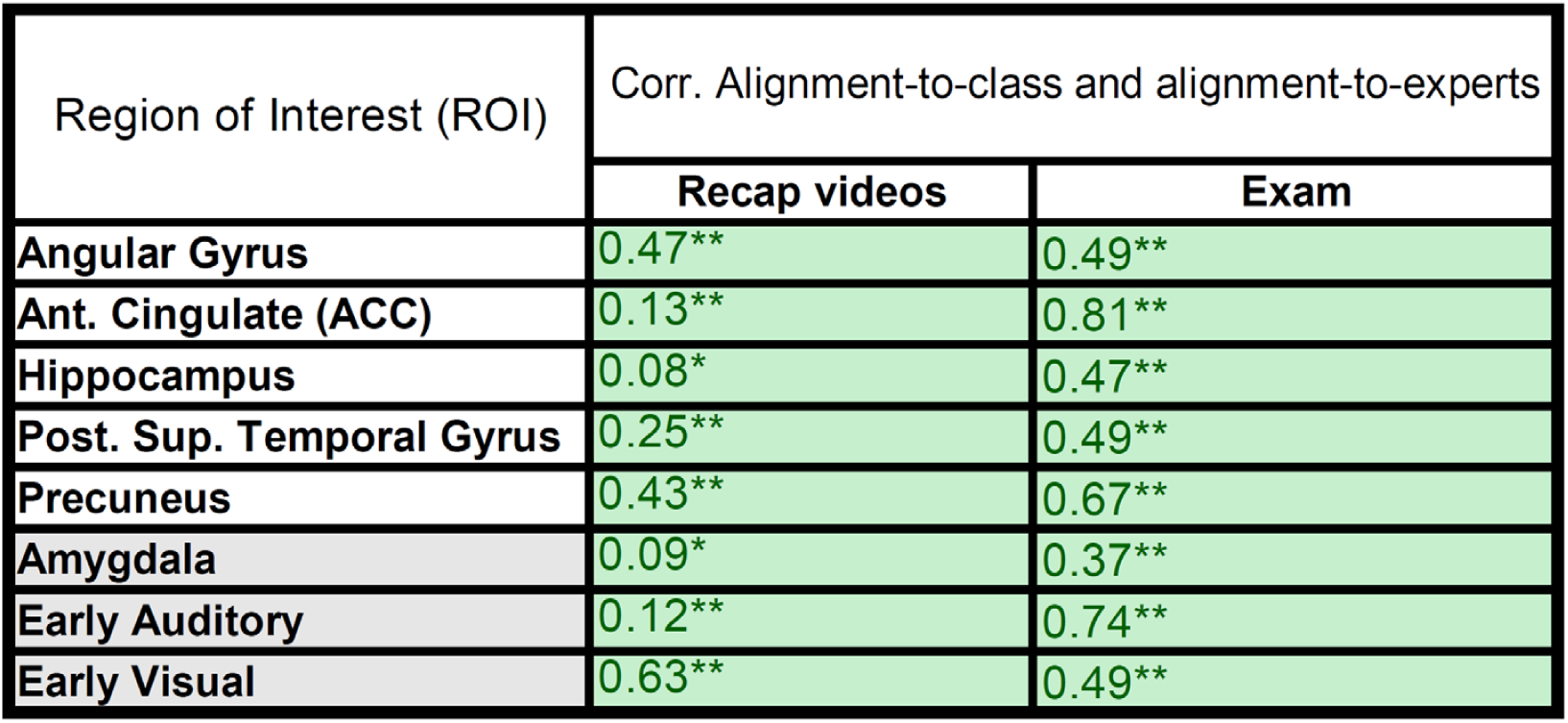
Alignment-to-experts is positively correlated with alignment-to-class during recaps and during final exam. Correlation between alignment-to-class and alignment-to-experts is shown during lectures and during the exam. Results are shown in DMN ROIs as well as in control regions in sensory cortex (visual, intracalcarine cortex; auditory, Heschl’s gyrus) and in subcortex (amygdala). Green, significant correlation (permutation test, p<0.05, FDR corrected across 7 ROIs). n.s., not significant.

**Figure 3.**
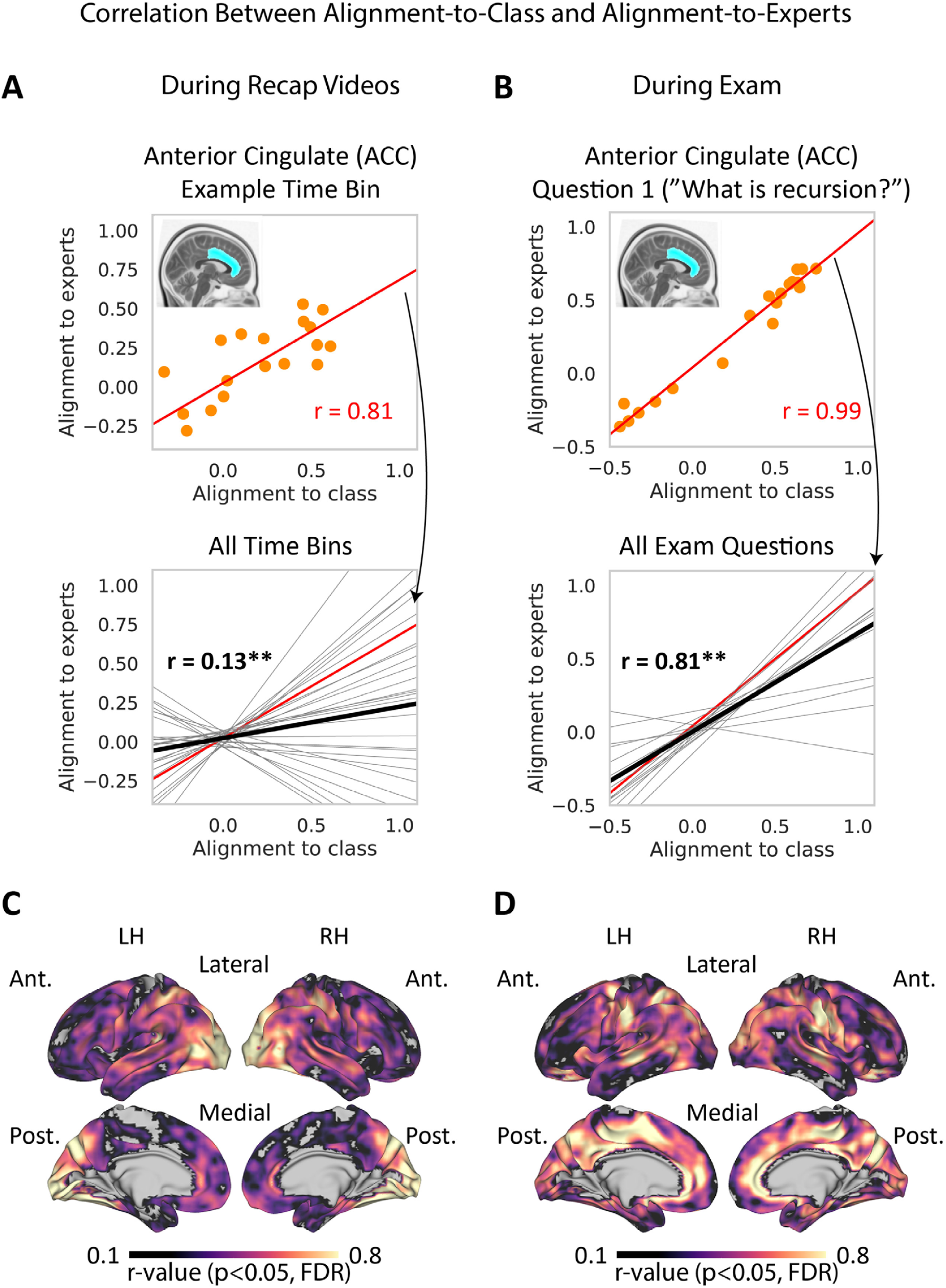
Alignment-to-class and alignment-to-experts are positively correlated across the brain. Correlation between alignment-to-class and alignment-to-expert during recap videos (left) and during final exam (right) are shown. **A**. Between-subjects correlation during recap videos, in a single ROI. Top, correlation in a single 30-second time bin. Orange dots represent individual students. Bottom, mean across all time bins (solid black line). Trendlines for individual time-bins are shown in grey, with the example time bin shown in red. **B**. Between-subjects correlation during the final exam, in a single ROI. Top, correlation during the first question. Orange dots represent individual students. Bottom, mean across all exam questions (solid black line). Trendlines for individual questions are shown in grey, with the example question shown in red. See table 3 for a summary of ROI analysis results. **C**. Correlation during recap videos across the cortex, searchlight analysis results shown. **D**. Correlation during the final exam across the cortex. Voxels showing significant correlation are shown in color. LH, left hemisphere, RH, right hemisphere, Ant., anterior, Post., posterior.

### Think like an expert: assessing understanding during exam using expert canonical responses

We hypothesized that better alignment to experts and to peers during question answering would be linked to better answers. To test this hypothesis, we obtained spatial activity patterns during each question and calculated “same-question” alignment-to-experts and alignment-to-class scores (Fig. 4A, see Methods for details). These scores allowed us to quantify how the neural patterns evoked by each question were related to the neural patterns evoked by the same question in other participants. We correlated alignment and question scores separately (across questions) within each student (see Fig. 4B for an example from a single student in a single ROI), and then took the mean across all students (Fig. 4C). Importantly, this within-participant design allowed us to capitalize on between-questions variability while controlling for individual differences.

**Figure 4.**
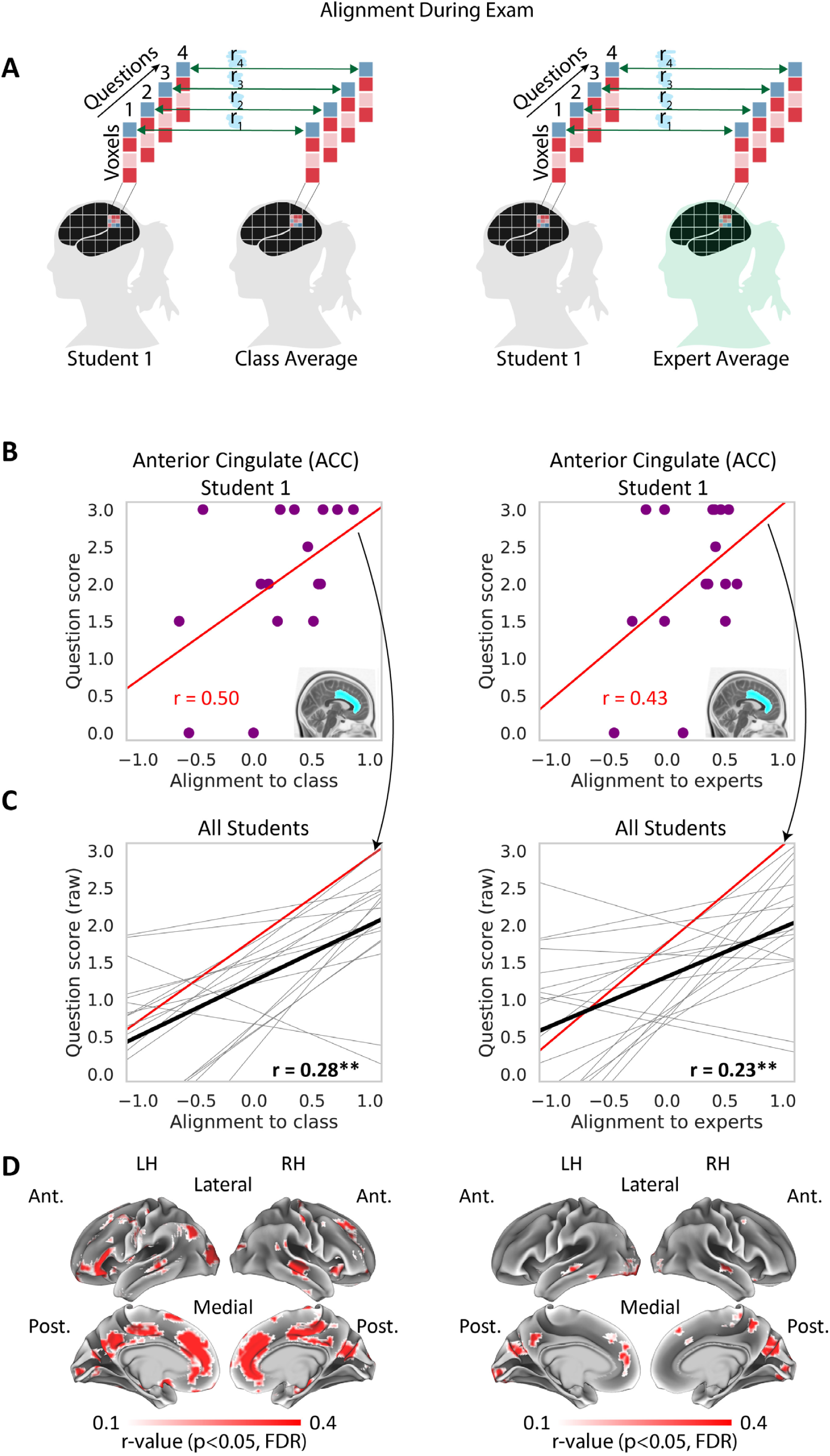
Same-question alignment during the exam correlates with performance. **A**. Left, student and class patterns are correlated on a question-by-question basis to derive alignment-to-class during exam. Right, student and expert patterns are similarly correlated to derive alignment-to-experts. **B**. Within-subject correlation between alignment and exam score in a single ROI, in a single student. Violet dots represent individual exam questions. Left, correlation between alignment-to-class and exam score. Right, correlation between alignment-to-experts and exam score. **C**. Within-subject correlation between alignment and exam score in a single ROI, trendlines for all students shown. Red, the trendline of the student shown in panel B. Black, mean across all students. Left, correlation between alignment-to-class and exam score. Right, correlation between alignment-to-experts and exam score. For all ROI analysis results, see table 3. **D**. Correlation across the cortex, searchlight analysis results shown. Voxels showing significant correlation are shown in color. Left, correlation between alignment-to-class and exam score. Right, correlation between alignment-to-experts and exam score. Control analyses for response length are shown in Fig. S2A and S2B. Note the correspondence between the two maps in major DMN nodes on the medial surface. LH, left hemisphere, RH, right hemisphere, Ant., anterior, Post., posterior.

Alignment-to-experts and alignment-to-class were both positively correlated with exam scores across several ROIs. In the ACC and superior temporal ROIs, alignment-to-class and alignment-to-experts were both positively correlated with exam scores (Fig. 4D, Table 2). Exam scores were also significantly correlated with alignment-to-experts in the precuneus and alignment-to-class in the hippocampus, angular gyrus and visual ROIs. Our searchlight analysis results supported these findings, highlighting regions across anterior and posterior medial cortex bilaterally (Fig. 4D). Importantly, both alignment-to-experts and alignment-to-class searchlight results highlighted these medial cortical regions. These findings show that neural alignment of specific question-by-question patterns was associated with better learning outcomes, indicating that concepts that were represented more similarly to the experts (and the class) were the concepts that students better understood. The results further highlight the ACC and medial prefrontal (mPFC) regions as areas where both alignment-to-class and alignment-to-experts were significantly correlated with behavior.

We then turned to examine the link between neural alignment and behavior while controlling for response length. This was motivated by the possibility that (i) longer answers might have yielded more stable spatial patterns, and that (ii) response length and quality could be linked (e.g. better answers could be longer). Therefore, a possible alternative explanation for our results is that they were driven by response length. To address this, we used a within-participant regression model to predict question scores from answer length. This model yielded a residual error term for each question (“residual score”, predicted score minus true score). We then repeated our original analysis using the “residual score” instead of the true score for each question. This procedure yielded cortical maps that were highly similar to those shown here (Fig. S2 A,B). Thus, across all brain areas showing a link between alignment and exam performance, effects were robust to response length.

### “Knowledge Structure” reflects learning in individual students

In the next set of analyses, we asked how individual concepts were integrated together in learners’ brains. Specifically, we hypothesized that learning new concepts also entails learning their contextual relations to other concepts. For example, the concepts “binary tree” and “linked list” are related in a specific way (a linked list can be used to implement a binary tree). To test this, we first created a “knowledge structure” for each participant, capturing the set of relationships between neural patterns evoked by different questions. For each question in each participant, we measured the similarity of the neural pattern evoked by that question to the canonical patterns evoked by *other* questions (in the class average or in the experts). That is the knowledge structure for that question for that participant. To predict performance, we then compared that question-specific knowledge structure (for that participant) to the question-specific knowledge-structure for the experts (alignment-to-experts) or for the class as a whole (alignment-to-class) (Fig. 5A). The resulting alignment scores were then correlated with question scores using a within-participant design (Fig. 5B).

**Figure 5.**
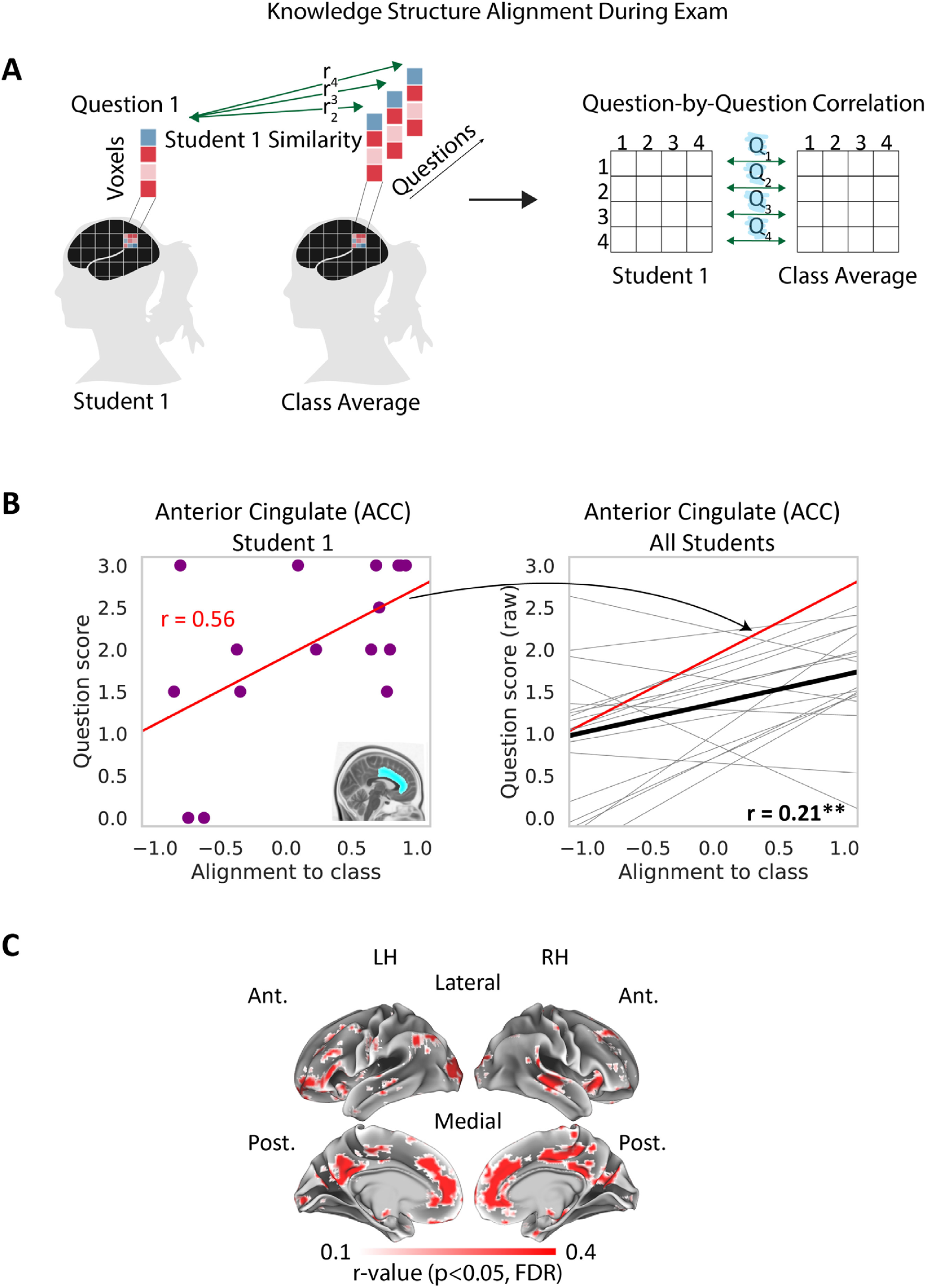
“Knowledge Structure” alignment during the exam correlates with performance. **A**. Left, student and class knowledge structures are correlated on a question-by-question basis to derive knowledge structure alignment during exam. Cell i,j in the student’s knowledge structure is the correlation between the student’s pattern for question i with the class pattern for question j (left). Student and mean class knowledge structures are then correlated on a row-by-row (question-by-question) basis. **B**. Within-subject correlation between alignment-to-class and exam score in a single ROI, in a single student. Each violet dot represents a single question. Left, correlation between alignment-to-class and exam score. Right, within-subject correlation between alignment-to-class and exam score in a single ROI, trendlines for all students shown. Red, the trendline of the student shown in panel B. Black, mean across all students. **C**. Correlation between knowledge structure alignment-to-class and exam score, searchlight analysis results shown. Voxels showing significant correlation are shown in color. Left, correlation. A control analysis for response length is shown in Fig. S2C. Searchlight results for alignment-to-experts are shown in Fig. S3. Note the correspondence between the alignment-to-class maps here and in Fig. 4. LH, left hemisphere, RH, right hemisphere, Ant., anterior, Post., posterior.

We found that “knowledge structure alignment” was positively correlated with exam scores across the hippocampus, ACC, angular gyrus and temporal ROIs when derived for the student cohort (alignment-to-class, Table 2). In line with this, our searchlight analysis showed robust results for alignment-to-class, highlighting medial cortical regions (Fig. 5C). Furthermore, the searchlight analysis showed a remarkable correspondence between knowledge structure and same-question results, with both maps highlighting similar medial regions (Fig. 4D, 5C). ROI and searchlight analysis results for alignment-to-experts were not significant across all regions (p>0.05, corrected). While alignment-to-experts searchlight results were qualitatively similar to alignment-to-class results (Fig. S3, p<0.01, uncorrected), no voxels survived multiple comparisons correction. In sum, these results showed that (i) students’ exam performance was significantly tied to their ability to create - and reinstate - a specific set of relationships between neural representations; and that (ii) the anatomical regions involved in knowledge structure alignment showed high correspondence with regions involved in same-question alignment. As a control, we repeated this analysis while controlling for response length, again obtaining highly similar results (Fig. S2C).

### Effects in DMN regions across tasks

Across our dataset, we repeatedly observed a link between learning outcomes and neural alignment in medial prefrontal regions, posterior medial regions, left angular gyrus, and medial temporal gyrus. We therefore performed an intersection analysis to substantiate this observation and determine whether the same, or different, voxels in these regions emerged across tasks. This analysis highlighted voxel clusters in anterior medial cortex, as well as in posterior medial cortex and superior temporal cortex, that showed significant effects across all alignment-to-class analyses (Fig. 6A). This set of regions overlaps in large part with the DMN. Furthermore, the intersection of the correlation map of same-question alignment-to-experts with exam scores and the correlation map of same-question alignment-to-class with exam scores yielded a similar map (Fig. 6B). These results indicated a key role for DMN regions across different phases of learning and further emphasized the link between alignment-to-experts and alignment-to-class measures.

**Figure 6.**
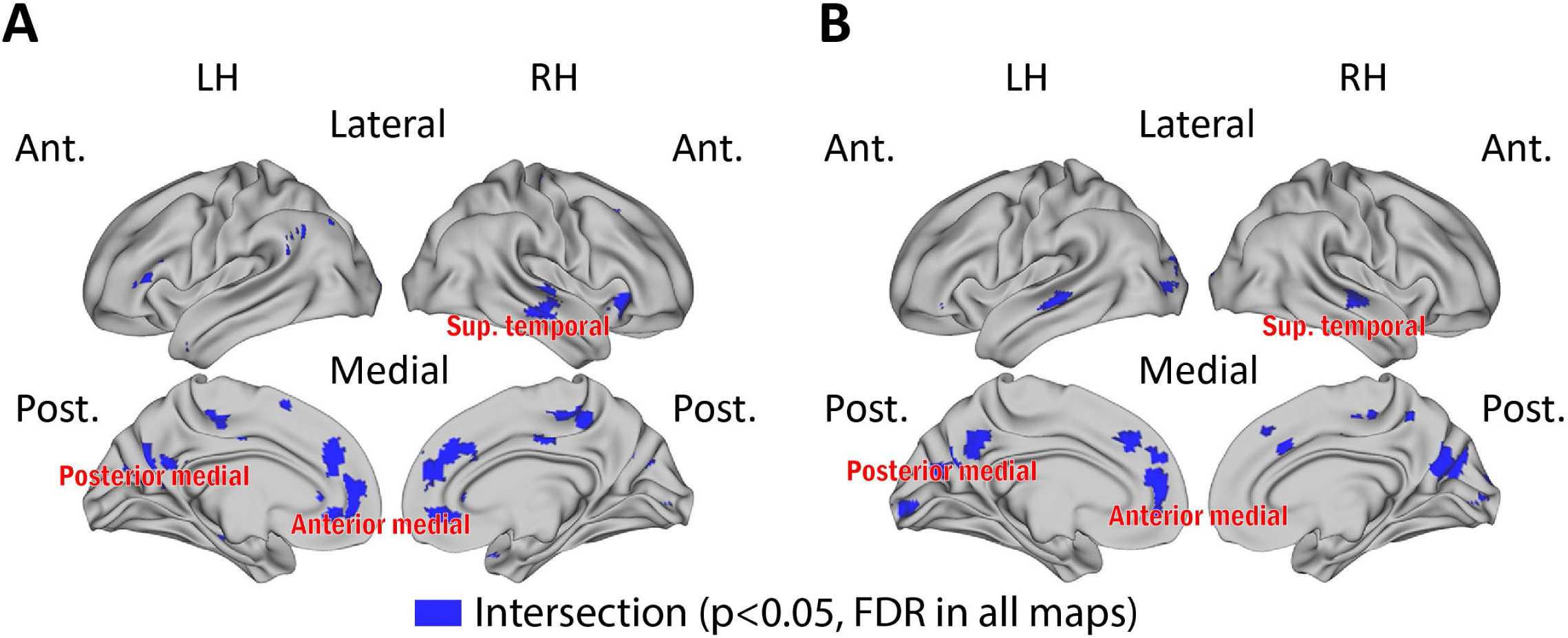
Robust neural alignment effects in medial and temporal cortical regions emerge across all analyses. **A**. Overlap regions across all three datasets and analyses for alignment-to class. Blue color indicates voxels in the intersection set of the following maps: (i) correlation between alignment-to-class during lectures and exam scores (shown in Fig. 2D), (ii) correlation between alignment-to-class and alignment-to-experts during recaps (shown in Fig. 3C), (iii) correlation between same-question alignment-to-class during the final exam and exam score (shown in Fig. 4D, left panel), and (iv) correlation between knowledge structure alignment-to-class during the exam and exam score (shown in Fig. 5C). **B**. Overlap regions for same-question analyses, blue color indicates voxels in the intersection set of the following maps: (i) correlation between same-question alignment-to-class during the final exam and exam score (shown in Fig. 4D, left panel), (ii) correlation between same-question alignment-to-experts during the final exam and exam score (shown in Fig. 4D, right panel).

## Discussion

The rapid changes in the field of education and the recent push towards online learning triggered by the recent pandemic have highlighted the need for novel teaching and assessment tools. In this work, we introduce a neural approach to predicting and assessing learning outcomes in real-life settings. This approach hinges on the idea that successful learning involves forming the “right” neural representations, which are captured in canonical activity patterns shared across learners and experts. In the current study, we put forward the notion that understanding is mirrored in “neural alignment”: the degree to which individual learners’ neural representations match canonical representations observed in experts. We tested this hypothesis in students enrolled in an introduction to computer science course and in graduate student “experts”, using a longitudinal fMRI design. Our findings show that across regions involved in memory encoding and reinstatement in the DMN and hippocampus, alignment successfully predicted overall student performance in a final exam. Furthermore, within individual students, concepts that evoked better alignment were better understood. We discuss the role of neural alignment in learning and understanding below.

### Neural alignment successfully predicts learning outcomes

During learning, neural activity patterns in each student participant comprised both common and idiosyncratic components. This is in line with a growing body of work showing that neural alignment across individuals watching the same video or listening to the same audio narrative is positively correlated with the level of shared context-dependent understanding (Chen et al., 2017; Nguyen et al., 2019; Yeshurun et al., 2017; Zadbood et al., 2017). Our results show that alignment-to-class was strongly correlated with exam score: across students, stronger similarity to the class predicted better performance (Fig. 2). This observation held across the Default Mode Network, implicated in internally focused thought and memory (Buckner et al., 2008; Hassabis and Maguire, 2007; Rugg and Vilberg, 2013). These results dovetail with findings that better alignment to common patterns in these regions supports better memory for shared experiences (Chen et al., 2017; Zadbood et al., 2017). The current results are also in line with recent EEG findings linking higher temporal synchrony (inter-subject correlation, ISC) during short educational videos with higher motivation and better learning outcomes (Cohen et al., 2018; Zhu et al., 2019). Importantly, our results extend this line of work to a real-world college course setting, allowing us to directly assess understanding and predict student performance.

A key point here is that, in our “flipped” class, a significant part of learning occurred outside of lectures. Given the structure of the course, it is unlikely that alignment-to-class during any specific lecture directly reflected the understanding of lecture topics at the end of the course. The first viewing of a course lecture, like the first reading of a textbook chapter, is just the beginning of a learning process that includes repetition and practice. Furthermore, only a fraction of course lecture segments (∼1/7 of total) was shown to student participants in the scanner, while performance was measured using an exam deliberately designed to span the entire course. Therefore, to explain the predictive power of alignment-to-class, we need to go beyond lecture-specific effects. We submit that neural alignment to common patterns reflects the online, moment-to-moment process of learning within individuals. Furthemore, the results indicate that monitoring such a process can predict to some extent the outcome of the learning process. This claim is supported by the finding that it is possible to reliably predict learning outcomes from neural activity during the early weeks of the course (Fig. S1). Other evidence pointing in the same direction can be found in an imaging study that linked children’s math IQ score with neural synchrony with adults during brief math videos (Cantlon and Li, 2013).

### Class patterns reflect expert patterns

To understand why alignment with the class leads to improved performance, we need to consider what shared class patterns may reflect. One possibility is that these patterns reflect group understanding. According to this view, when individual patterns are averaged and idiosyncratic differences cancel out, what emerges is a good approximation of an ideal “canonical” representation. Thus, the mean is a reflection of the fact that most students, most of the time, follow the lecture as intended: what they share is the correct interpretation of course material. A caveat is that common misunderstandings would also be reflected in the common signal. These misunderstandings, however, would not be shared by experts. Thus, if shared class patterns are “canonical”, we should expect them to match expert patterns. We tested this hypothesis during recaps (short summaries of the lecture videos, shown just prior to the exam) and during the exam. Our findings confirmed that alignment-to-class and alignment-to-experts were positively correlated across large swaths of the cerebral cortex, including in DMN regions (Fig. 3). The tight link between alignment-to-class and alignment-to-experts suggests that students and experts may converge on a single set of shared neural states.

### Alignment tracks understanding of specific topics

In direct support of our alignment-as-understanding hypothesis, we found that alignment and understanding were correlated on a fine-grained, question-by-question basis, within individual students. Our results show that during the exam, alignment-to-experts and alignment-to-class were both positively correlated with question scores (Fig. 4). Importantly, these results were specific to the neural patterns observed in the experts for each particular question and thus were robust to individual differences. In other words, our results could not emerge due to some students being better learners than others, or having better working memory, for example. The effects tested here could only emerge if, in individual students, answers that evoked better alignment-to-experts obtained higher scores and vice versa.

To our knowledge, this is the first demonstration of shared structure in neural responses across individuals during open question answering. Formulating an answer required participants to call upon their memory and understanding of question-specific concepts, as well as more general cognitive processes such as language production. The correlation of alignment and performance emerged most strongly in medial DMN regions, suggesting that the aligned neural patterns in these areas supported introspection and memory (figure 6B). It is therefore possible that successful alignment reflected understanding, particularly in light of the body of work linking similarity in DMN regions to better understanding of narratives (Chen et al., 2017; Zadbood et al., 2017). This opens up the future possibility of using alignment to assess understanding, offering a different perspective than traditional performance measures.

### Learning the right “knowledge structure”

The more abstract a concept, the less it is grounded in physical reality. This has posed a challenge to teachers, who need to build a structure of interrelated ideas from the ground up. In a course like “Introduction to Computer Science”, the understanding of basic concepts (e.g. “algorithms”) later facilitates the introduction of more advanced theoretical concepts (e.g. “intractability”). The alignment of knowledge structures across students provides a fine-grained measure of understanding specific topics. It shows how each topic is grounded in others, revealing the interaction between mental representations. This result could therefore allow examining understanding in individual learners in high resolution. While same-question alignment-to-class could show, for example, that the concept of “intractability” was not well understood, knowledge structure alignment could show that the underlying reason is difficulty with the more basic concept of “recursion”. Our knowledge structure analysis results further suggest that, even at the end of the course, students’ knowledge structure did not converge on that of experts. We speculate that, unlike specific topic knowledge, experts’ knowledge structure draws more heavily on their broader understanding of the field. Another possibility is that this null result is due to lack of power (fewer experts than students in our dataset). In sum, these results show that the set of relationships between mental representations of abstract concepts is behaviorally relevant, and point to medial DMN regions as key nodes supporting these representations. Future work will focus on delineating the exact relationship between student and expert knowledge structures.

### A key role for medial DMN regions during learning

The significance of DMN cortical structures in our results is in line with previous work that localized behaviorally-relevant, memory-related shared representations to these areas (Chen et al., 2017; Zadbood et al., 2017). They are also consistent with earlier findings that specific patterns of activity during memory encoding in DMN regions predicted recall performance (Bird et al., 2015), as well as with a recent report of hippocampal changes triggered by learning the structures and names of organic compounds (Just and Keller, 2019). The posterior medial (PM) cortical system plays a key role in episodic memory (Ranganath and Ritchey, 2012) as well as forming part of the DMN. However, studies that examined the neuronal correlates of math and science have generally highlighted cortical areas outside the DMN. For example, parietal and frontal regions have been shown to play a key role in mathematical cognition (Anderson et al., 2011; Dehaene et al., 2004). In physics, concepts such as gravity and frequency have each been associated with a distinctive set of cortical regions, mostly on the lateral cortical surface (Mason and Just, 2016), and recent work using multivariate methods has localized representations of physics concepts to dorsal fronto-parietal regions and ventral visual areas (Cetron et al., 2019). One way to account for these apparent discrepancies and for the prominence of DMN regions in our outcome-based results is to consider the likely role of different cortical regions in learning. While specific types of cognitive operations may well be subserved by distinct sets of cortical regions, long-term learning requires forming the correct neural representations, encoding them in memory and retrieving them in the right context, all hallmarks of the DMN and its associated structures.

An unexpected finding is that, during lectures, we also observed correlations between exam scores and alignment-to-class in early sensory areas (Fig. 2, Table 2). A possible explanation for such correlations during lectures is that these correlations reflected top-down effects that interacted with the way students processed visual and auditory information (for example, it is possible that stronger learners attended to specific details in the lectures video which were missed by less attentive students).

### Limitations

Despite the wealth of fMRI data in this study, our results were derived from scanning a cohort of students enrolled in a single course at a single campus. Further research is required in order to ensure that they generalize to other domains and learning settings. Although we see no a-priori reason why our findings should be limited to any particular type of course (e.g. courses in STEM, or introductory courses), or a particular type of college, further research is required to rule out these possibilities.

## Conclusions

In this work, we tracked neural activity patterns in individual learners as they were taking a demanding STEM course. We found that neural activity converged on “canonical” patterns shared across learners and with experts. The degree of alignment to canonical patterns tracked understanding of individual highly abstract topics in individual students, and predicted success in a final exam. While a wide application of neural measures outside the lab would require finding a suitable alternative to MRI scanning (or working around its limitations), measuring learners’ alignment to canonical neural representations has the power to potentially transform online learning and its typical one-lesson-fits-all approach. How best to achieve these goals remains a topic for future research.

## Methods

### Participants and stimuli

Twenty-four “student” and five “expert” participants (11 female) were recruited for the study. All participants were right-handed, had normal or corrected-to-normal vision and hearing, and reported no learning disabilities. All except one expert were native English speakers. Student participants reported having no prior knowledge or experience in computer science. Prior to scanning, all students completed the course placement exam (described under “Stimuli”) in written form and received 0 out of 3 points on all questions (see grading details below). Experts all had an undergraduate or graduate degree in computer science and reported significant knowledge in the field. Participants received monetary compensation for their time. All participants provided informed written consent in accordance with experimental procedures approved by the Princeton University IRB.

Students were enrolled in COS 126: Computer Science - An Interdisciplinary Approach (lectures available at informit.com/title/9780134493831) and were taking the course for the first time. The course sets out to teach basic principles of computer science in the context of scientific, engineering, and commercial applications. It uses a “flipped” classroom model, with students viewing lecture videos on their own schedule and interacting with course staff in precepts and class meetings. All students took the course for credit and participated in the course normally, with the exception that they were asked to view part of the lecture videos (∼3 hours out of ∼21 hours in total) in the scanner. Subjects were asked not to view these lecture videos online before the scans. Students were scanned every 2-3 weeks during a single semester (Fig. 1). Four students dropped the course and were excluded from the experiment. Two student datasets were incomplete (one student skipped scan 3; one student’s exam scan data was not collected due to experimenter error). One expert did not complete the exam. The final sample consisted of twenty datasets collected from undergraduate students (18 complete) and five expert datasets (four complete). No statistical methods were used to predetermine sample sizes. Our student sample size is similar to those reported in previous publications from our group (Chen et al., 2017; Honey et al., 2012; Regev et al., 2013).

Stimuli included video lectures, recaps and a final exam, shown throughout a series of six scans. During each of the first five scans, students watched 3-5 segments of course lecture videos that were required viewing for the following week (mean segment length 9 minutes, total of ∼40 minutes shown in each scan, total of 21 segments in all scans). At the end of each scan, students were given a set of questions about the lecture (question data not analyzed in the current manuscript). In addition, on scans 3-5, students were shown two 3-minute recap videos, summarizing the previous two lectures shown (this data was not analyzed in the current manuscript). On the final scan, students watched all five 3-minute recap videos, each summarizing a single lecture (“recaps”). This was followed by an exam that required verbal responses. The same stimuli were shown to all students. Experts underwent the final scan only. Each lecture segment and recap was shown in a separate scanner run. At the beginning and end of each run, we appended 20-30 seconds of unrelated “filler” audiovisual clips (from YouTube “oddly satisfying” compilations, featuring, for example, objects being assembled neatly). Filler clips were similarly added in previous studies from our group (Chen et al., 2017; Nguyen et al., 2019). This was done because the stimulus onset may elicit a global arousal response, which could add noise to the analysis. To avoid this, scan data collected during fillers, as well as during the first 12 seconds of each video, were omitted from analysis. Exam stimuli consisted of 16 written questions, shown in fixed order. We used the course placement exam, developed by course staff for the benefit of students wishing to demonstrate proficiency in course material without taking it. Questions were designed to span the breadth of material covered in the course; some required distilling large concepts into simple explanations and others were more practical. Exam scores were not used by course staff to assess students’ performance. The same exam was used to assess students’ knowledge prior to scanning (in written form) and on the final scan (with verbal responses).

### Experimental procedures

Participants were asked to watch lecture videos as they normally would. Lecture videos were shown at normal speed (first scan) or slightly accelerated (x1.15 speed, scans 2-5). Stimuli were projected using an LCD projector onto a rear-projection screen located in the scanner bore and viewed with an angled mirror. PsychoPy2 was used to display the stimuli and synchronize them with MRI data acquisition (Peirce et al., 2019). Audio was delivered via in-ear headphones (Sensimetrics S14), and the volume was adjusted for every participant before each scan. Video monitoring was used to monitor participants’ alertness in about 40% of scans at random (Eyelink, SR Research). Monitoring showed that no participants fell asleep during the experiment. Verbal responses to exam questions were recorded using a customized MR-compatible recording system (FOMRI III, OptoAcoustics). Motion during speech was minimized by instructing participants to remain still and by stabilizing participants’ heads with foam padding, as in previous studies from our group (Nguyen et al., 2019; Simony et al., 2016; Zadbood et al., 2017). Participants indicated end-of-answer using a handheld response box (Current Designs).

No outside resources were available during the videos or the exam, and students could not take notes. No feedback was provided to participants during the exam. The exam was self-paced with a “Please Wait” text slide presented for 12 seconds between questions. Question text was shown for the entire length of the answer at the center of the screen, and participants confirmed they could read it easily. Participants could start giving a verbal answer ten seconds after question onset, indicated by the appearance of a countdown clock at the bottom of the screen (90 seconds per question; no time-outs were recorded). Data collected between questions and during the first 8 seconds of each question were truncated to avoid including non-question responses. Verbal responses to exam questions were anonymized and transcribed by two of the authors (MM and HH) using open source software (Audacity, www.audacityteam.org). Transcripts were then scored by two independent raters (teaching assistants on the course staff) on a scale of 0-3, and the mean was taken. Written exams taken by the students prior to scanning were rated in a similar way. Expert responses rated below 2 were omitted from analyses (6 out of 64 responses in total). This was done to ensure expert brain activity patterns reflected correct answers. No student responses were omitted. Students’ total exam score (sum of all 16 questions) was normalized to a standard 0-100 scale.

### fMRI acquisition

MRI data were collected on two 3-T full-body scanners (Siemens Skyra and Prisma) with 64 channel head coils. Scanner-participant pairing was kept constant throughout the experiment. Functional images were acquired using a T2*-weighted echo-planar imaging (EPI) pulse sequence (TR 2000 ms, TE 28 ms, flip angle 80 deg, FOV 192 × 192 mm^2^, whole-brain coverage with 38 transverse slices, 3 mm^3^ voxels, no gap, GRAPPA iPAT 2). Anatomical images were acquired using a T1-weighted MPRAGE pulse sequence (1 mm^3^ resolution).

### fMRI preprocessing

Preprocessing was performed in FSL 6.0.1 (http://fsl.fmrib.ox.ac.uk/fsl), including slice time correction, motion correction, linear detrending, high-pass filtering (100 s cutoff) and gaussian smoothing (6mm FWHM) (Jenkinson et al., 2002, 2012). Functional volumes were then coregistred and affine transformed to a template brain (Montreal Neurological Institute). Motion parameters (3 translations and 3 rotations) were regressed out from functional data using linear regression. All calculations were performed in volume space. Data were analyzed using Python 3 (www.python.org) and R (www.r-project.org), using the Brain Imaging Analysis Kit (http://brainiak.org) and custom code. Eight regions of interest (ROIs) were anatomically defined using the probabilistic Harvard-Oxford cortical and subcortical structural atlases (Desikan et al., 2006). ROIs were defined across major Default Mode Network (DMN) nodes in the angular gyrus, precuneus, and anterior cingulate cortex (ACC), as well as in the hippocampus and posterior superior temporal gyrus, and control regions in early visual cortex (intracalcarine sulcus), early auditory cortex (Heschl’s gyrus) and in sub-cortex (amygdala). Bilateral ROIs were created by taking the union of voxels in both hemispheres. A liberal threshold of >20% probability was used. To avoid circularity, all voxels within the anatomical mask were included and no functional data was used to define ROIs. Projections onto a cortical surface for visualization purposes were performed, as a final step, with Connectome Workbench (Marcus et al., 2011).

### Alignment during lectures

Multi-voxel BOLD patterns during lectures were obtained as follows. First, we used 30-second non-overlapping bins to extract multi-voxel activity (table 1). This yielded a single pattern for every bin in every participant. Then, to examine spatial similarities between participants during videos, we employed an inter-subject pattern correlation framework, which has been successfully used to uncover shared memory-related responses (Chen et al., 2017; Nastase et al., 2019; Zadbood et al., 2017). For each pattern in each student, we obtained an alignment-to-class measure by directly comparing the student pattern and the mean class pattern (average across all other students), using Pearson correlation. Then, correlation values were averaged within video segments. Throughout the manuscript, correlation values were transformed by Fisher’s *z* prior to averaging and then back-transformed in order to minimize bias (Silver and Dunlap, 1987). Finally, we averaged across segments to obtain a single alignment-to-class measure for every student during all lectures. Alignment was derived independently for each ROI and each searchlight. We then used alignment to predict student performance in the placement exam. To this end, we used a between-participants design, correlating alignment and overall exam scores (mean across questions). Statistical significance values were derived using a one-sided permutation test, with a null distribution created for each searchlight by shuffling score labels 1000 times.

### Power analysis across lectures

We defined a “stable prediction index” across the cortex by considering the effect of information accumulation throughout the lectures on prediction success. To this end, we used the alignment-to-class values calculated for each one of our 21 individual lecture segments. We started by correlating exam scores with alignment-to-class in the first video (scan 1, segment 1). We then proceeded in sequence, correlating exam scores with the mean of alignment-to-class values across segments 1 and 2, and finally with the mean across all lecture segments. We performed this process for every cortical voxel using searchlight, to obtain a series of 21 r-values for each voxel (one for each added segment). As before, a p-value was calculated for each r-value using a one-sided permutation test by randomizing score labels. Using a liberal threshold of p<0.01 (uncorrected), we considered all voxels that showed a significant correlation between exam scores and alignment-to-class as calculated above. Lowering the threshold allowed us to include all potentially predictive voxels. For each voxel, we then defined the “stable prediction index” as the number of segments required to (a) reach a significant correlation between exam score and alignment and (b) maintain significance for all subsequently added segments (no “breaks”). By design, a high index number (21) showed that data from all lecture segments was required to achieve a significant correlation and thus reflected late prediction. In contrast, a low index number (1) showed that significant prediction could be obtained by considering data from the first lecture segment alone, affording early prediction of exam score. Index values were calculated independently for each ROI and each searchlight.

### Correlation between alignment-to-class and alignment-to-experts

Alignment-to-class during recaps was derived similarly to lectures. For each 30-second time bin in each student, we obtained an alignment-to-class measure by directly comparing the student pattern and the mean class pattern (average across all other students), using Pearson correlation. In addition, we obtained an alignment-to-experts measure by comparing the student pattern and the mean pattern across experts (“canonical” pattern). We thus obtained an alignment-to-class and alignment-to-experts pattern for each student, in each bin. We used Pearson correlation to correlate these alignment measures using a between-participants design, obtaining a single correlation value in each time bin. Finally, we took the mean across all time bins within each recap, and then across recaps. Statistical significance values were derived using a one-sided permutation test, with a null distribution created for each time bin by shuffling student labels 1000 times. Correlation between alignment-to-class and alignment-to-experts during the exam was performed in an analogous manner, with questions used in place of time bins (see below).

### Neural alignment to experts during the exam

Students’ and experts’ multi-voxel activity patterns during the exam were obtained by taking the mean fMRI BOLD signal during each question, in each participant. Each spatial pattern thus reflected neural responses associated with the specific subset of course topics included in that question. To compare student and class patterns, we again used the inter-subject pattern correlation framework. We derived an alignment-to-class score by correlating each question pattern in each student with the class average of the same question (mean across all other students) and then taking the mean across all students (Fig. 4A). We performed this on a question-by-question basis to obtain a vector of 16 alignment-to-class values for each student (one value for each question). Similarly, we derived an alignment-to-experts score by correlating each question pattern in each student with the mean pattern across experts, and obtained a vector of 16 alignment-to-expert values (one value for each question). Alignment was derived independently for each ROI and each searchlight. We then correlated alignment and question scores within students using Pearson correlation, obtaining a single r-value for each student. Finally, we took the mean across students. Statistical significance values were derived using a one-sided permutation test. We created a null distribution for each student by shuffling score labels 1000 times and then compared the mean across students to the mean null distribution.

### “Knowledge structure” alignment

We defined “knowledge structures” as similarity matrices aimed at capturing the set of relationships between question representations. In the following, we describe how a student-specific knowledge structure was constructed and correlated with a class-derived template and an expert-derived template to derive (i) alignment-to-class and (ii) alignment-to-expert scores. Finally, we describe how within-participant correlation was used to examine the link between alignment and performance in individual students. First, we used the canonical class average and expert average patterns calculated for each question (see above) and constructed two templates. A class template was constructed by correlating “canonical class” question patterns with each other. This yielded a 16 question x 16 question symmetric similarity matrix comprising the distances between pairs of question patterns (r-values) (figure 5A). Similarly, an expert template was constructed by correlating “canonical expert” question patterns with each other. This yielded two 16 question x 16 question symmetric similarity matrices comprising the distances between pairs of question patterns (r-values). We then constructed a “knowledge structure” matrix for every student by correlating each question pattern (in that student) with the template pattern of all other questions. A single row in this structure thus represented the similarity between a student’s neural response to a specific question and the template (class/expert) responses to every other question. Next, we correlated student and template matrices, row by row, excluding the diagonal, and obtained a question-by-question alignment score for each student. For each student, we thus derived a vector of 16 alignment-to-class values (one value for each question), and a vector of 16 alignment-to-expert values. Lastly, we correlated alignment and question scores within students using Pearson correlation, obtaining a single r-value for each student, and took the mean across students. A null distribution was created for each student and a one-sided permutation test was used to determine statistical significance as before.

### Intersection analysis

Intersection maps across data-driven searchlights were created by examining statistically significant voxels across analyses (p<0.05, corrected). Figure 6A shows the intersection of the following maps: (i) correlation between alignment-to-class during lectures and exam scores (shown in Fig. 2D), (ii) correlation between alignment-to-class and alignment-to-experts during recaps (shown in Fig. 3C), (iii) correlation between same-question alignment-to-class during the final exam and exam score (shown in Fig. 4D, left panel), and (iv) correlation between knowledge structure alignment-to-class during the exam and exam score (shown in Fig. 5C). Figure 6B shows voxels in the intersection set of the following maps: (i) correlation between same-question alignment-to-class during the final exam and exam score (shown in Fig. 4D, left panel), (ii) correlation between same-question alignment-to-experts during the final exam and exam score (shown in Fig. 4D, right panel).

## Author Contributions

M.M., L.H., H.H., K.A.N and U.H. designed the experiment. M.M., L.H., H.H., Y-F.L. and M.N. collected the data. M.M., K.A.N and U.H. analyzed the data and wrote the manuscript.

## Acknowledgments

The authors wish to thank Princeton COS 126 staff and in particular Robert Sedgewick, Dan Leyzberg, Christopher Moretti, Kevin Wayne, Ibrahim Albluwi, Bridger Hahn, Thomas Schaffner, and Rachel Protacio; Mona Fixdal and the McGraw Center for Teaching and Learning; The Scully Center for the Neuroscience of Mind and Behavior; Peter J. Ramadge; and members of the Norman and Hasson labs for fruitful discussions. This study was supported by NIH Grant DP1-HD091948 to U.H. and by Intel Labs.

## Resources availability

The data that supports the findings of this study will be made available online. Analysis code and exam questions used in the study are available from the corresponding author upon request.

## Supplementary Information

### Supplementary Results

#### Prediction of learning outcomes prediction improves with number of lecture segments

We performed a power analysis across lectures to determine the amount of neural data required in order to obtain robust correlations with exam scores (i.e. how early in the course we could predict learning outcomes). This was motivated by our desire to inform future studies and applications of our measures to real-world scenarios, where resource optimization (i.e. less scanning) may be desired. To this end, we first obtained alignment-to-class values for each student in each lecture segment (21 in total). Then, we correlated exam scores with alignment in the first segment, the first two segments, and so forth until information from all segments was accumulated. This has allowed us to examine changes in score prediction quality due to the accumulation of information across lectures. An ROI analysis showed that, in the hippocampus, prediction quality increased steadily as more data was added, and afforded significant prediction after a single scan (Fig. S1). To test whether this was the case across the cortex, we calculated a “Stable Prediction Index” for every voxel in the brain using searchlight (see Methods). On this index, a low number corresponded to regions where few data points were required to achieve significant correlation with behavior (i.e. early prediction), and a higher number to regions where more data points were required (i.e. late prediction). We found significant variance within and across cortical regions. Thus, while alignment-to-class in some parts of the angular gyrus afforded early prediction, only late prediction was possible in other parts (i.e. given the entire dataset).

#### Supplementary Figures

**Figure S1.**
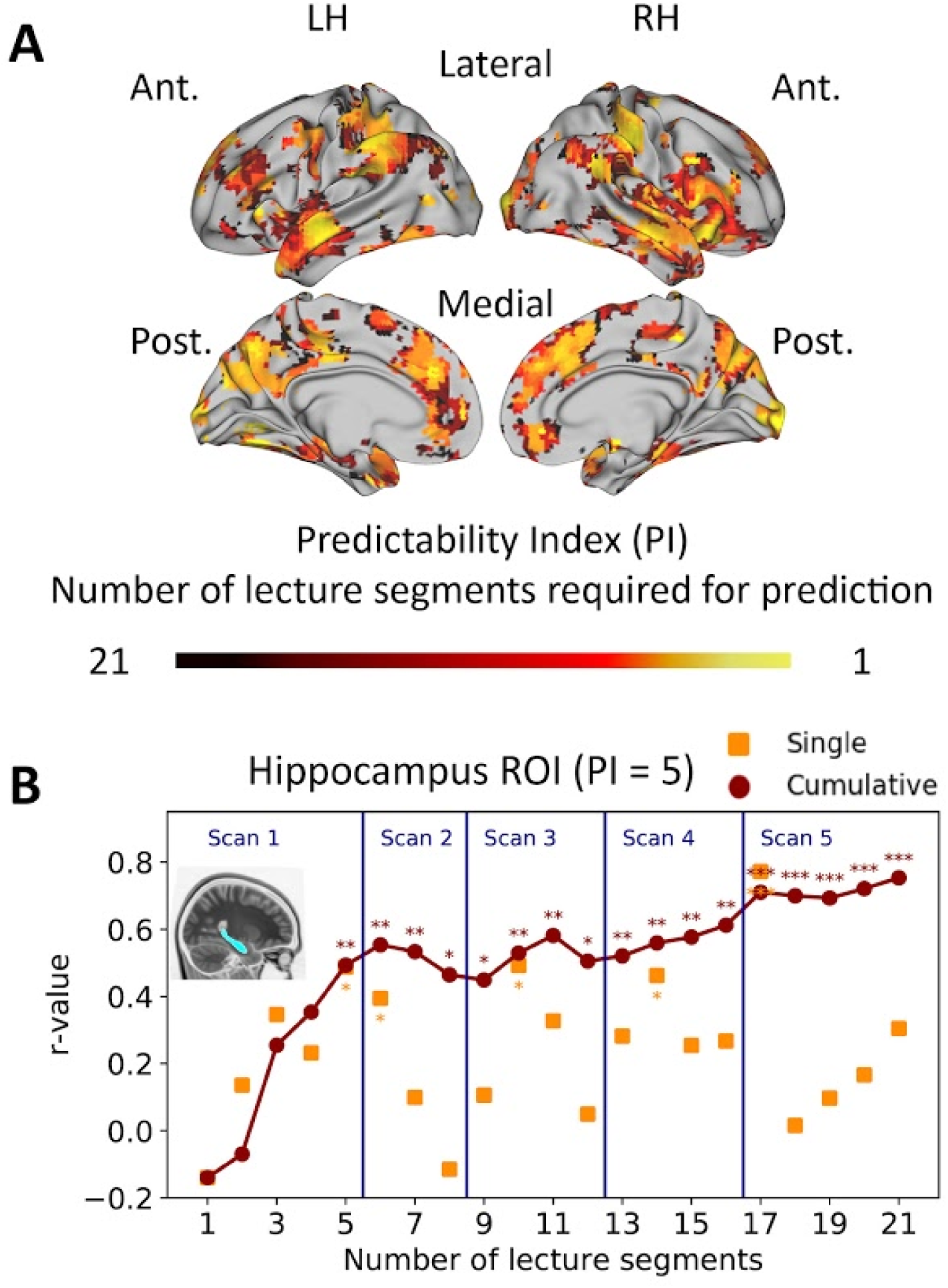
Variance across the brain in the number of lectures required for performance prediction. **A**. Searchlight analysis results. Per-voxel “predictability index” values shown. Note the low index scores across major DMN nodes, indicating prediction of exam score can be achieved with a small number of lecture segments. **B**. Number of lectures required for stable prediction in the hippocampus. Yellow rectangles, prediction result for individual lecture segments (correlation between exam scores and alignment-to-class in that segment). Brown line, prediction of exam score from data accumulated over lecture segments.

**Figure S2.**
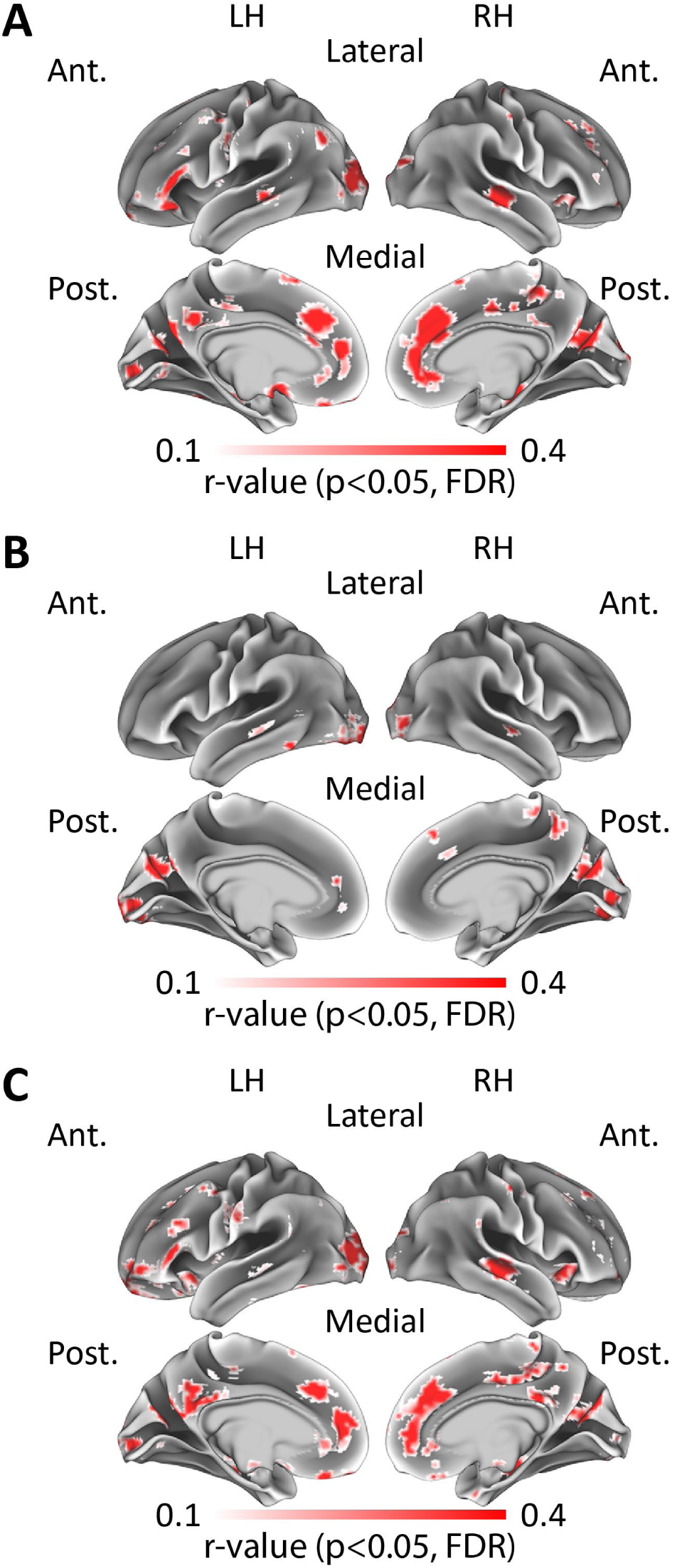
“Same-question” and “Knowledge structure” effects controlled for response length. Searchlight analysis results shown. Voxels showing significant correlation are shown in color. **A**. Correlation between same-question alignment-to-class and exam score, controlled for response length. **B**. Correlation between same-question alignment-to-experts and exam score, controlled for response length. **C**. Correlation between “knowledge structure” alignment-to-class and exam score, controlled for response length. Note the close correspondence between these maps and the results of the original analyses in Fig. 4D and Fig.5C. LH, left hemisphere, RH, right hemisphere, Ant., anterior, Post., posterior.

**Figure S3.**
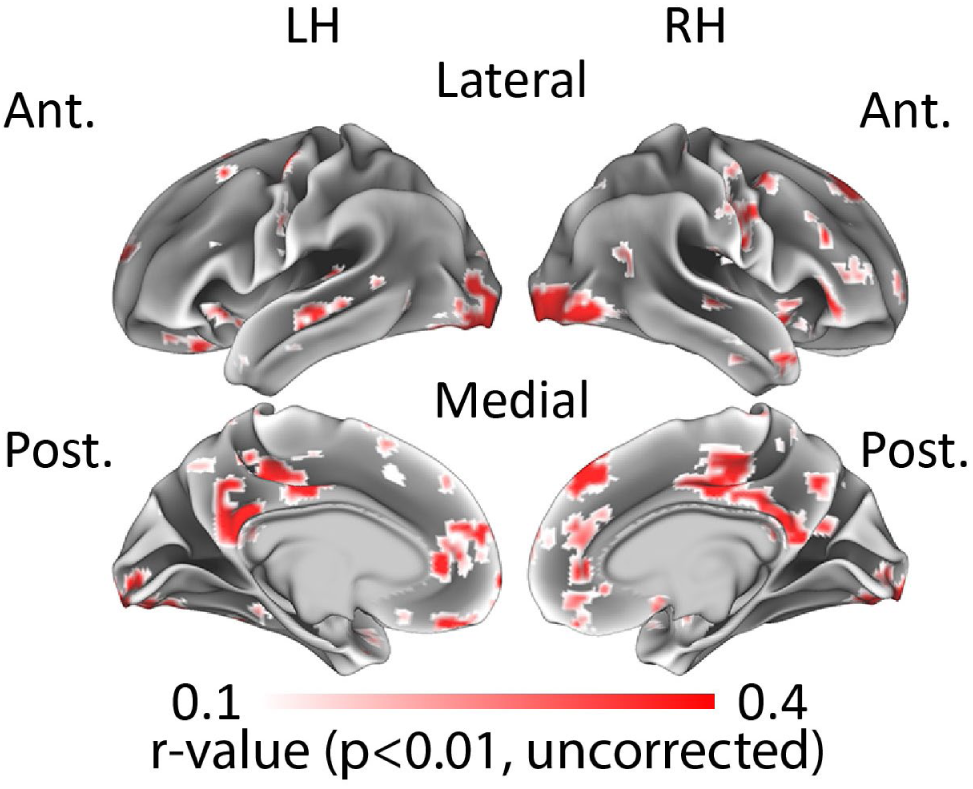
Correlation of “knowledge structure” alignment-to-experts and performance. Searchlight analysis results shown. Map thresholded using a liberal statistical threshold (p<0.01, uncorrected). No voxels survived multiple comparisons correction (p<0.05, FDR). Note the qualitative similarities to knowledge structure alignment-to-class results in medial cortical structures (Fig. 5C). LH, left hemisphere, RH, right hemisphere, Ant., anterior, Post., posterior.

